# Light-sheet engineering using the Field Synthesis theorem

**DOI:** 10.1101/700633

**Authors:** Bo-Jui Chang, Reto Fiolka

**Affiliations:** Department of Cell Biology, UT Southwestern Medical Center, Dallas, TX, USA; Lyda Hill Department of Bioinformatics, UT Southwestern Medical Center, Dallas, TX, USA

**Keywords:** Field Synthesis, lattice light-sheet, C-light-sheet

## Abstract

Recent advances in light-sheet microscopy have enabled sensitive imaging with high spatiotemporal resolution. However, the creation of thin light-sheets for high axial resolution is challenging, as the thickness of the sheet, field of view and confinement of the excitation need to be carefully balanced. Some of the thinnest light-sheets created so far have found little practical use as they excite too much out-of-focus fluorescence. In contrast, the most commonly used lightsheet for subcellular imaging, the square lattice, has excellent excitation confinement at the cost of lower axial resolving power. Here we leverage the recently discovered Field Synthesis theorem to create light-sheets where thickness and illumination confinement can be continuously tuned. Explicitly, we scan a line beam across a portion of an annulus mask on the back focal plane of the illumination objective, which we call it C-light-sheets. We experimentally characterize these light-sheets and their application on biological samples.

## Introduction

Light-sheet fluorescence microscopy (LSFM) has been transformative for volumetric imaging of single cells up to entire organisms, as it minimizes sample irradiation and allows efficient and rapid 3D imaging^1–3^. LSFM provides excellent spatiotemporal resolution, but is commonly not considered a super-resolution technique, as its spatial resolving power is still diffraction limited. Nevertheless, some LSFM implementations have been developed that can attain 300nm scale axial resolution^3–7^, effectively doubling the resolution of confocal microscopy, the workhorse in 3D microscopy. This is enabled by the larger set of angles that the separate illumination and detection objectives cover compared to a single objective microscope system. As such LSFM can improve the z-resolution over epi-fluorescence microscopes without compromising temporal resolution or requiring specialized fluorophores or non-linear optical phenomena.

A driver for z-resolution in light-sheet microscopy is the thickness of the sheet, which in turn is governed by the laws of diffraction. For Gaussian beams, the thickness of the sheet is coupled via beam divergence to the confocal parameter of the beam waist, which dictates over what propagation distance a light-sheet can be approximated. While in principle Gaussian light-sheets with sub-micron thickness can be created, their confocal parameter is usually shorter than a typical cell, which makes them ill-suited for practical imaging.

Propagation invariant beams, such as Bessel^8^ and Airy^9, 10^ beams, and optical lattices, can in principle overcome the divergence of Gaussian beams. To create a light-sheet, such a beam is rapidly scanned laterally, a process known as digitally scanned light-sheet (DSLM)^2^. However, when propagation invariant beams are used for DSLM, for a given light-sheet thickness, any increase in confocal parameter is paid by reduced confinement of the excitation energy into the sheet. This has been first discovered with Bessel beam light-sheets^11–14^: while the main lobe of such a light-sheet is very narrow, it is accompanied by a large beam skirt that may contain over 90% of the energy within the light-sheet^6^. Thus large amounts of out-of-focus light are excited, which generate unwanted out-of-focus fluorescence and lead to accelerated photo-bleaching.

To address this problem, the Betzig lab introduced lattice light-sheet microscopy (LLSM)^3^, which generates a light-sheet by coherent superposition of many Bessel beams. By carefully adjusting the spacing between the individual beams, optical lattices can be generated that can balance the thickness and the confinement of the light-sheet. The most widely used sheet in LLSM is the square lattice, which offers excellent confinement, but consists of a much thicker main lobe than a Bessel beam light-sheet. For higher axial resolution, the hexagonal lattice was proposed, which has a thinner main lobe than the square lattice, however at the cost of much stronger sidelobes. So far hexagonal lattice light-sheets have been rarely used, presumably because the sidelobes are difficult to remove numerically.

Recently, we have discovered a new mathematical theorem called Field Synthesis that can be leveraged to create light-sheets in a more flexible and general way^15^. In Field Synthesis, a line is scanned over a mask conjugate to the back pupil of the illumination objective and a light-sheet is generated as an incoherent summation of the resulting instantaneous intensity distributions (see also Figure 1). While we have shown previously that this method can be used to recreate existing light-sheets faithfully, we explore here the potential of Field Synthesis to generate new light-sheets that are tailored to specific applications. In particular, our goal was to create light-sheets that have properties in between hexagonal and square lattices. Here we present experimental results of these new light-sheets that we have termed C-sheets, which are characterized in transmission and with fluorescent nanospheres, and are applied to biological imaging.

**Figure 1.**
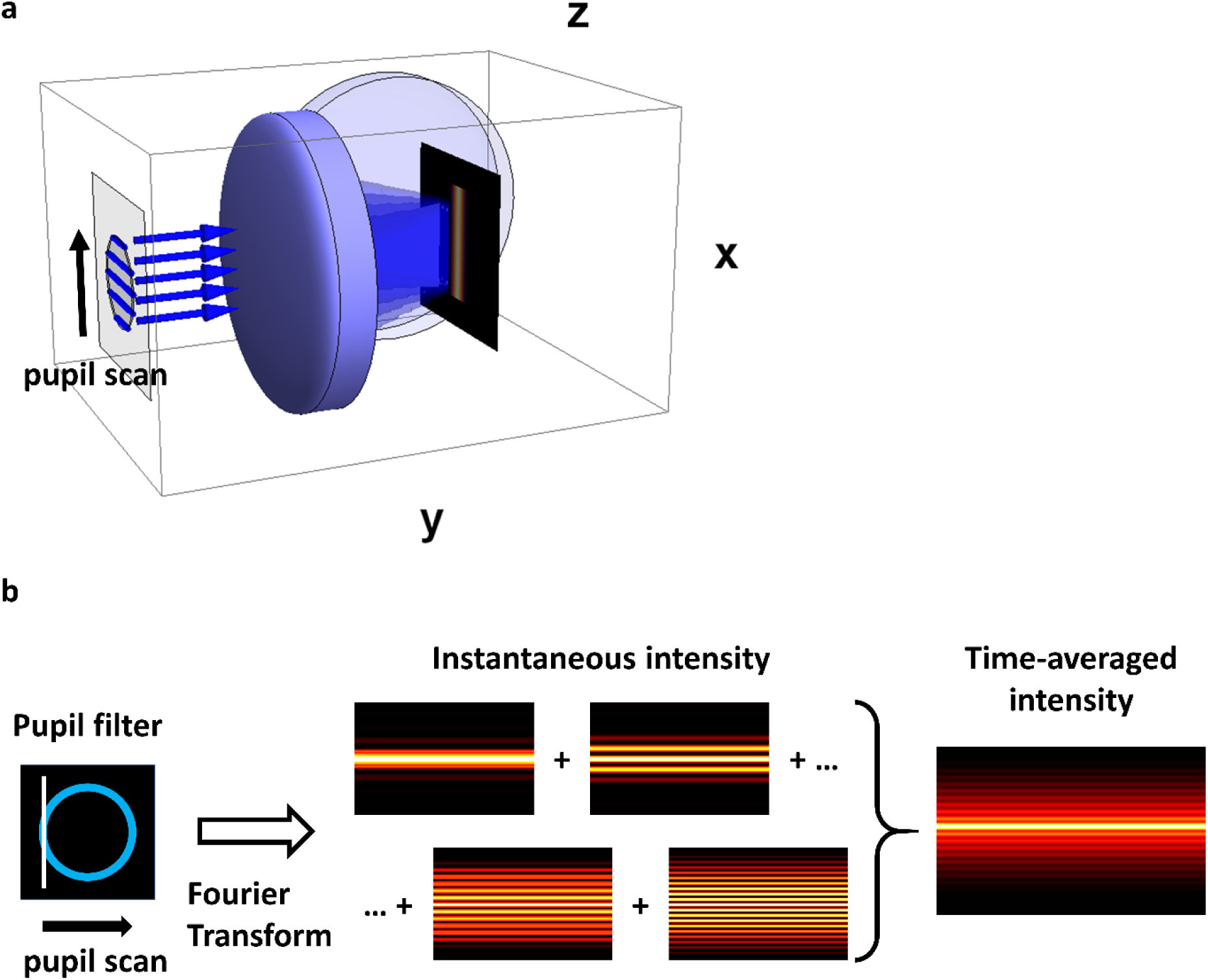
**a**. Schematic sketch of Field Synthesis. A line is scanned over a pupil mask that is conjugate to the back pupil of an objective. Five blue lines are shown to symbolize five positions of the scan. **b**. Field Synthesis of a Bessel beam light-sheet. A line is scanned over an annulus in the pupil plane. For each scan position, an instantaneous intensity pattern is formed in real space. By summing up all these patterns, a Bessel beam light-sheet emerges.

### Concept: Light-sheet design

For the studies here, we restricted ourselves to an annular shaped pupil mask. The design rationales are outlined for light-sheet generation using the Field Synthesis approach (Figure 1), where a line is scanned over a pupil filter in Fourier space. As shown in Figure 2a, when analyzing the image formation process for one particular scan position of this line, the Fourier components of the light-sheet are given by the autocorrelation of the pupil function, in this case resulting in three line segments. These components give rise in real space to intensity distributions as shown in figure 1b, a sinusoidal pattern with a compact support. We note that a purely sinusoidal pattern would only result for an infinitely thin annulus mask. Such illumination pattern have recently been described as Bessel beams^16, 17^, however, mathematically they are Cosine patterns bound by an envelope. For simplicity, we will call such a beam a Cosine-Gauss beam.

**Figure 2.**
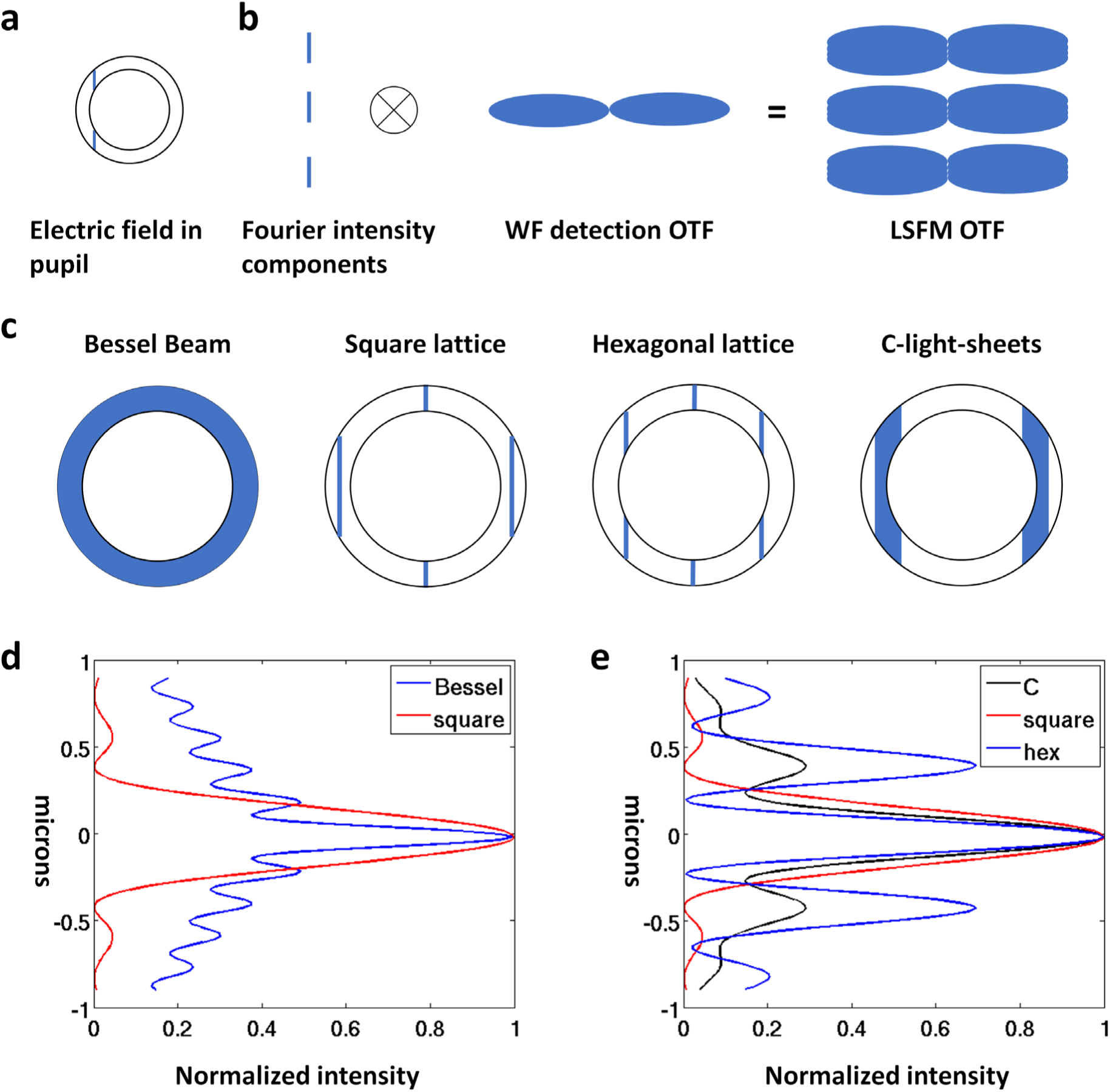
Light-sheets design. **a.** Electric field in the pupil of the illumination objective for one scan position in Field Synthesis. **b.** Simplified sketch for image formation in Field Synthesis for the scan position shown in a. WF: Wide-field, OTF: Optical transfer function. LSFM: light-sheet fluorescence microscopy.^®^ denotes the convolution operation **c.** Pupil functions for Bessel, square and hexagonal lattices, and C-light-sheets. Blue color indicates transmitted laser light through the annular pupil filter. **d.** Comparison of axial profiles of a Bessel beam and a square lattice light-sheet. **e.** Comparisons of a C-light-sheet, square lattice light-sheet and hexagonal lattice light-sheet. d. and e. are simulation results.

Using such a Cosine-Gauss beam as a light-sheet, we can compute the overall LSFM optical transfer function (OTF), which is the convolution of the Fourier components of the illumination and the wide-field optical transfer function (Figure 2b). As it can be seen, there are gaps between the main three lobes of the LSFM OTF. This has been previously described in standing wave microscopy^18^, where a sinusoidal illumination pattern is created by interfering two laser beams from two opposing objectives. A remedy was readily found by Lanni and later Gustafsson et al by using multiple standing waves of varying axial frequencies to fill the gaps^19, 20^.

This can be done for light-sheet illumination as well. If we scan a line over the full annulus (Figure 2c), a continuum of Cosine-Gauss beams is superimposed that contain the lowest spatial frequency (edges of the annulus) up to the highest spatial frequency that the optical system can produce (center of the annulus). As we have previously shown, the resulting time-averaged intensity distribution corresponds to a Bessel beam light-sheet^15^ (Figure 2d).

If we instead use Field Synthesis to create a square lattice light-sheet, the beam is stepped to three discrete positions in the pupil, two at the edges of the annulus and one at its center (Figure 2c). The resulting light-sheet has significantly reduced ringing, but its main lobe is thicker than a Bessel beam light-sheet.

For a hexagonal lattice the two main orders are shifted more inwards to the center of the annulus, and hence create a light-sheet with higher spatial frequency content. Through simulations we found that the central order is much weaker and does not contribute significantly to the final light-sheet (supplementary note). Thus, in an approximation, one can describe the light-sheet as a single Cosine-Gauss beam. Such a light-sheet features a thin main lobe, at the cost of stronger sidelobes compared to a square lattice sheet (Figure 2e). Imaging with strong sidelobes has been shown to be a challenge in other imaging modalities, such as 4pi^21^ and I5M^20^, and the removal of the resulting ghost images is difficult^22^. As a practical rule of thumb, the sidelobe strength should be kept below 50% (for a theoretical derivation, see Nagorni and Hell^23^). This is the main reason that 4pi microscopy has only been widely used with two-photon excitation, which suppresses the sidelobes to a tolerable level^24^.

We hypothesized that we could create light-sheets with advantageous properties if we superimposed the spatial frequency range that exists between a square and a hexagonal lattice light-sheet. A continuum of intermediate frequencies can be generated if we scan a line from the position of the main order of the square lattice to the corresponding position of the hexagonal lattice (see Figure 2c). We expected that this would lead to a reduction in sidelobe strength (Figure 2e) and also would fill gaps in the OTF. We note that there is some similarity of this approach to light-sheets using Mathieu beams^25,26^ However in our case, the light-sheets are not generated by scanning a confined beam laterally, but by summing multiple Cosine-Gauss beams, resulting in a higher spatial duty cycle that reduces photo-bleaching^15^. Since the scanning pattern in the pupil looks like a letter C, we chose to call them C-light-sheets.

## Methods

### Optical setup

To create the light-sheets, we used the illumination train of our previously published Field Synthesis setup, but we used different objectives: we replaced the illumination lens with a special optics NA 0.69 lens and used a Nikon NA 1.1/25x lens for detection. 3D intensity images of the light-sheets were acquired in transmission, as previously described^15^. The annulus of the pupil filter corresponded to an inner NA of 0.45 and an outer NA of 0.536. To create the C-light-sheets, we scanned the beam across only one half of the annulus (see also Figure 3a for experimental results). Scanning both halves would not change the light-sheet itself but would add angular diversity that helps to suppress shadow artifacts. As of now, we have not implemented hardware control that would allow us to perform scanning on both sides of the annulus during one camera exposure.

**Figure 3.**
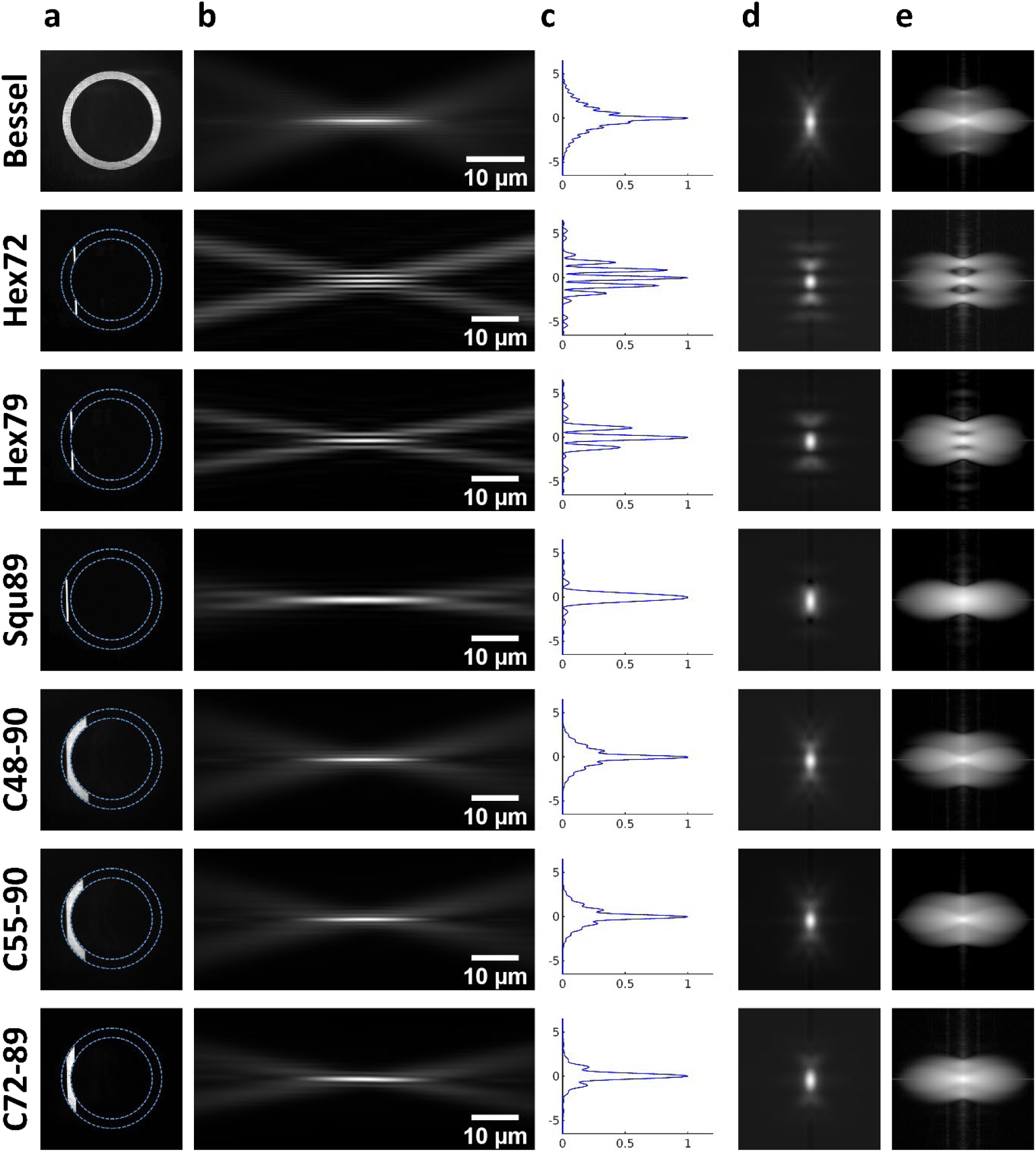
Measurements of light-sheets, point spread functions, and optical transfer functions. **a.** First column shows experimental images of the pupil plane. The numbers denote the beam position, normalized to the radius of the annulus. Hex stands for hexagonal lattice, Squ stands for square lattice, and C stands for C-light-sheet. For C-light-sheets, the numbers indicate the scan range, normalized to the radius of the annulus. **b.** Y-Z cross-sections through the different light sheets. The propagation direction of the light-sheets is from left to right. **c.** Axial (z) profiles of the light-sheets, which is obtained by the average of the light-sheet through about 40 microns field of view at the center of their beam waists. **d.** Experimental, rotationally averaged point spread functions for the different light-sheets. **e.** Optical transfer functions corresponding to the different light-sheets.

### Sample preparation

Bovine Type I collagen (Advanced Biomatrix, 5005-100) was labeled with AlexaFluor 488 NHS Ester (Fluoroprobes 1013-1) at 4 °C overnight in sodium bicarbonate buffer and dialyzed overnight at 4 °C in 0.2% acetic acid. Collagen samples were prepared by polymerizing 2 mg/ml of non-fluorescently labeled collagen supplemented with AlexaFluor488-labeled collagen in imaging holders. Briefly, 1 ml of non-fluorescent collagen solution was prepared by mixing 100 μl 10X phosphate buffered saline, 10 μl 1 N sodium hydroxide, 230 μl Milli-Q water and 660 μl of 3.0 mg/ml collagen stock solution. Next, 25 μl of AlexaFluor488-labeled collagen was added to 475 μl of the non-labeled collagen to achieve 5% labeling density. Collagen was allowed to polymerize at 37 °C in a humidified incubator, and stored in sterile phosphate buffered saline prior to imaging.

Native human retinal pigmented epithelium cells immortalized by HPV-16 (ATCC, CRL-2502) were used to generate stable cell lines overexpressing eGFP-Clathrin Light Chain (eGFP-CLCa, Loerke & Mettlen et al^27^) by retroviral infection. Cells were sorted for eGFP expression level (Moody Foundation Flow Cytometry, UTSW), and checked for incorporation of tagged CLCa. The cells used in these studies express 10 times more eGFP-CLCa than endogenous CLCa and exhibit a corresponding downregulation of endogenous to 10% as expected. eGFP-CLCa cells were cultured in DMEM/F12 supplemented with 10% FBS at 37 °C in a 5% CO2 atmosphere.

### Deconvolution

For each light-sheet 200 nm fluorescent nanospheres (Polysciences, 17151-10) were imaged and an isolated bead with a high signal to noise ratio was selected as a PSF, which was then rotationally averaged. The rotationally averaged PSF was subsequently refined by a blind deconvolution routine in MATLAB 2019a (deconvblind) over ten iterations. This helped to remove some small artifacts introduced by rotational averaging. We then used the retrieved PSF from the previous blind deconvolution to deconvolve image data (using again deconvblind). We limited the number of iterations to ten to avoid clipping of dim features and over-deconvolution (i.e. excessive deconvolution can produce overly optimistic resolution on point objects like beads).

## Results

We acquired the 3D intensity distribution of different light-sheets in transmission and acquired point spread functions using 200 nm fluorescent nanospheres. In Figure 3a-c, a comparison of different light-sheets is given. In the first column, the intensity as measured in a conjugate pupil plane is shown. The second column from the left (figure 3b) shows the different light-sheets as measured in transmission. Table 1 gives values for the beam waist and propagation length of the light-sheets.

**Table 1.** Light-sheet properties as measured in transmission. Mean and standard deviation are listed.

| Light-sheet type | Axial FWHM ($\mu\text{m}$) | Propagation FWHM ($\mu\text{m}$) | # of average |
| --- | --- | --- | --- |
| **Bessel** | 0.37±0.003 | 15.11±0.076 | 6 |
| **Hexagonal 72** | 0.30±0.001 | 13.69±0.215 | 6 |
| **Hexagonal 79** | 0.39±0.002 | 13.80±0.174 | 6 |
| **Square 89** | 0.69±0.010 | 22.19±0.607 | 6 |
| **Cshape 48-90** | 0.40±0.003 | 14.67±0.057 | 6 |
| **Cshape 55-90** | 0.42±0.003 | 14.72±0.098 | 6 |
| **Cshape 72-89** | 0.51±0.004 | 14.86±0.073 | 6 |

For a Bessel beam light-sheet (figure 3a, first row), a line is scanned over the full annulus. If we look at the light-sheet and its profile (figure 3c), a pronounced beam skirt is visible. This is reflected in the corresponding experimental point-spread function, which has a sharp peak at the center, but then gradually fades away. If we look at the corresponding OTF, one can see that the support is elongated in the axial direction, however the strength further away from the center is weak. This is a consequence of the strong excitation of out-of-focus light, which strengthens the lower frequencies in the LSFM OTF that correspond to the wide-field OTF.

In the next two rows, two hexagonal lattices are compared. Depending on the position of the line, the light-sheet exhibits sidelobes of varying strength (50% of the main lobe or higher). In the corresponding PSF, one can see that the main axial lobe can be squeezed, however, additional out-of-focus light is excited by the sidelobes. The corresponding OTFs feature two gaps in their support. This is a consequence of the strong sidelobes.

If we look at the square lattice light-sheet in the fourth row, it is obvious that it has very minimal sidelobes and thus excellent illumination confinement, however its PSF is much more elongated in the axial direction than in the case of the hexagonal lattices. Also, the OTF is very elliptical, meaning it has limited support in the axial direction.

In the last three rows, three different C-light-sheets are shown. In the corresponding axial profiles, one can see that the strength of the sidelobes is kept below 50% and that the intensity decays gradually without much ringing in the axial direction. The corresponding PSFs are more compact than the one of the square lattice, however they exhibit less out-of-focus blur than the hexagonal lattices. The optical transfer functions have no gaps and show a more solid support compared to the OTF of the Bessel beam.

Table 2 lists the lateral and axial full width half maxima (FWHM) of beads before and after deconvolution. In this measurement, 100 nm fluorescent nanospheres (Invitrogen, F8803) were used. The PSF of the Bessel light-sheets did not deconvolve well, and the axial resolution remains poor with ~800nm. In contrast, the square lattice light-sheet and the moderate C72-89 sheet showed little remaining sidelobes and improve the axial resolution to 577 and 478nm, respectively. Interestingly, the two C-sheets with a larger scan range (C48-90 and C55-90) did not improve axial resolution any further than the less aggressive C72-89.

**Table 2.** Lateral and axial resolution before and after deconvolution. Measured on 100nm fluorescent nanospheres, averaged over 10 spheres for each light-sheet. Mean and standard deviation are listed.

| Light-sheet type | Before deconvolution |  | After deconvolution |  |
| --- | --- | --- | --- | --- |
|  | Lateral FWHM (nm) | Axial FWHM (nm) | Lateral FWHM (nm) | Axial FWHM (nm) |
| **Bessel** | 295±14.2 | 848±45.5 | 282±11.2 | 818±37.0 |
| **Hexagonal 72** | 319±22.0 | 434±24.5 | 312±21.9 | 439±26.2 |
| **Hexagonal 79** | 313±22.4 | 532±21.8 | 257±16.7 | 421±13.2 |
| **Square 89** | 300±8.4 | 778±18.0 | 219±5.6 | 577±17.4 |
| **Cshape 48-90** | 303±14.4 | 660±36.5 | 266±11.7 | 550±24.3 |
| **Cshape 55-90** | 304±14.3 | 677±35.9 | 238±8.7 | 484±22.2 |
| **Cshape 72-89** | 305±9.5 | 661±14.7 | 228±6.8 | 478±9.1 |

While the hexagonal lattices can improve the axial resolution further (429-431nm), iterative deconvolution was not able to fully remove sidelobe structures, confirming the 50% sidelobe rule established for 4pi microscopy^23^. We hypothesize that the solid OTF support of the square and the C72-89 light-sheets help with deconvolution, especially in scenarios of low signal to noise ratios. Interestingly, this also held true for lateral resolution, where iterative deconvolution enabled a gain of about 1.36 for the square and the C72-89 light-sheets. For the other light-sheets, the gain in lateral resolution by deconvolution was notably smaller.

Next we volumetrically imaged collagen samples with Bessel, hexagonal and square lattice, and C72-89 light-sheets, followed by iterative deconvolution (Figure 4). Figure 4a shows a crosssectional view of collagen (maximum intensity projection) as imaged with the Bessel beam light-sheet. One can see that even after deconvolution, out-of-focus haze is still visible (inset on the left of figure 4a). In contrast, the square lattice light-sheet shows much higher optical sectioning and very little to no deconvolution artifacts (Figure 4b and inset on the right). While the hexagonal lattice light-sheet has higher resolving power in the axial direction, iterative deconvolution cannot completely remove some of the sidelobes (Figure 4c and inset on the left). In contrast, the C72-89 light-sheet enables artifact free deconvolution and resolves finer details than the square lattice light-sheet in the axial direction (Figure 4d and inset on the right).

**Figure 4.**
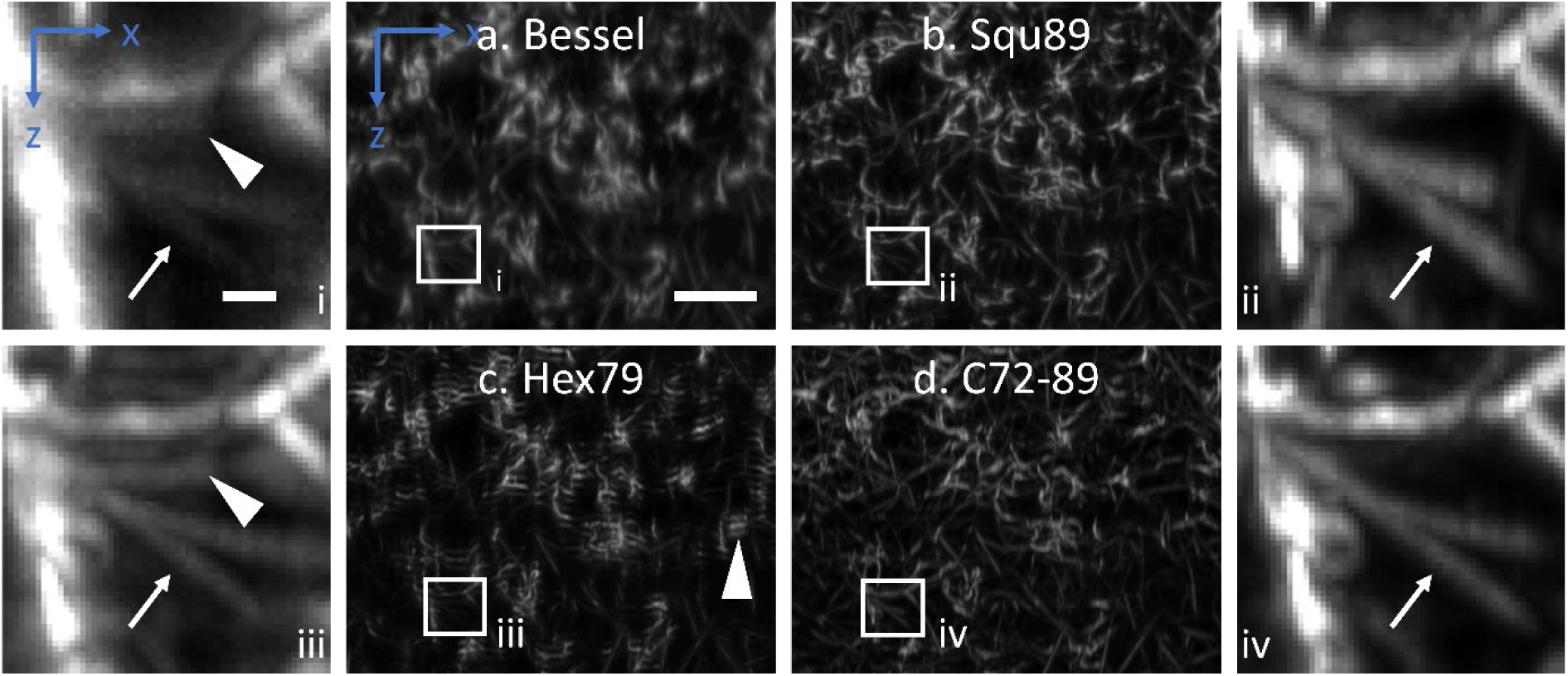
3D imaging of collagen using Bessel, square and hexagonal lattice, and C-light-sheets. **a.** Cross-sectional (x-z) maximum intensity projection image acquired with a Bessel beam light-sheet and followed by deconvolution. **b.** Same area as in a, as imaged by square lattice light-sheet followed by deconvolution. **c.** Same area as in a, as imaged by a hexagonal lattice light-sheet followed by deconvolution. **d.** Same area as in a, as imaged by a C72-89 light-sheet followed by deconvolution. Insets i-iv show magnified views of the boxed regions in a-d. Arrows mark two filaments, chevrons mark deconvolution artifacts. Scale bars: a: 10 microns; i: 1 micron.

We further imaged ARPE cells labeled with eGFP clathrin light-chain. Figure 5a shows maximum intensity projections (MIPs) of ARPE cells as imaged with Bessel, hexagonal and square lattice, and three different C-light-sheets. Figure 5b shows the magnified regions (red box in Figure 5a, single slice and MIPs of some planes), which highlights that the Bessel and hexagonal light-sheets do not deconvolve properly without artifacts. In contrast, the square lattice and the C72-89 light-sheet resulted in very cleanly deconvolved data. The more aggressive C-light-sheets (C55-90 and C48-90) show in individual slices (Figure 5b, bottom) slight blur above and below the main lobe of its PSF. In a maximum intensity projection, these artifacts are less visible.

**Figure 5.**
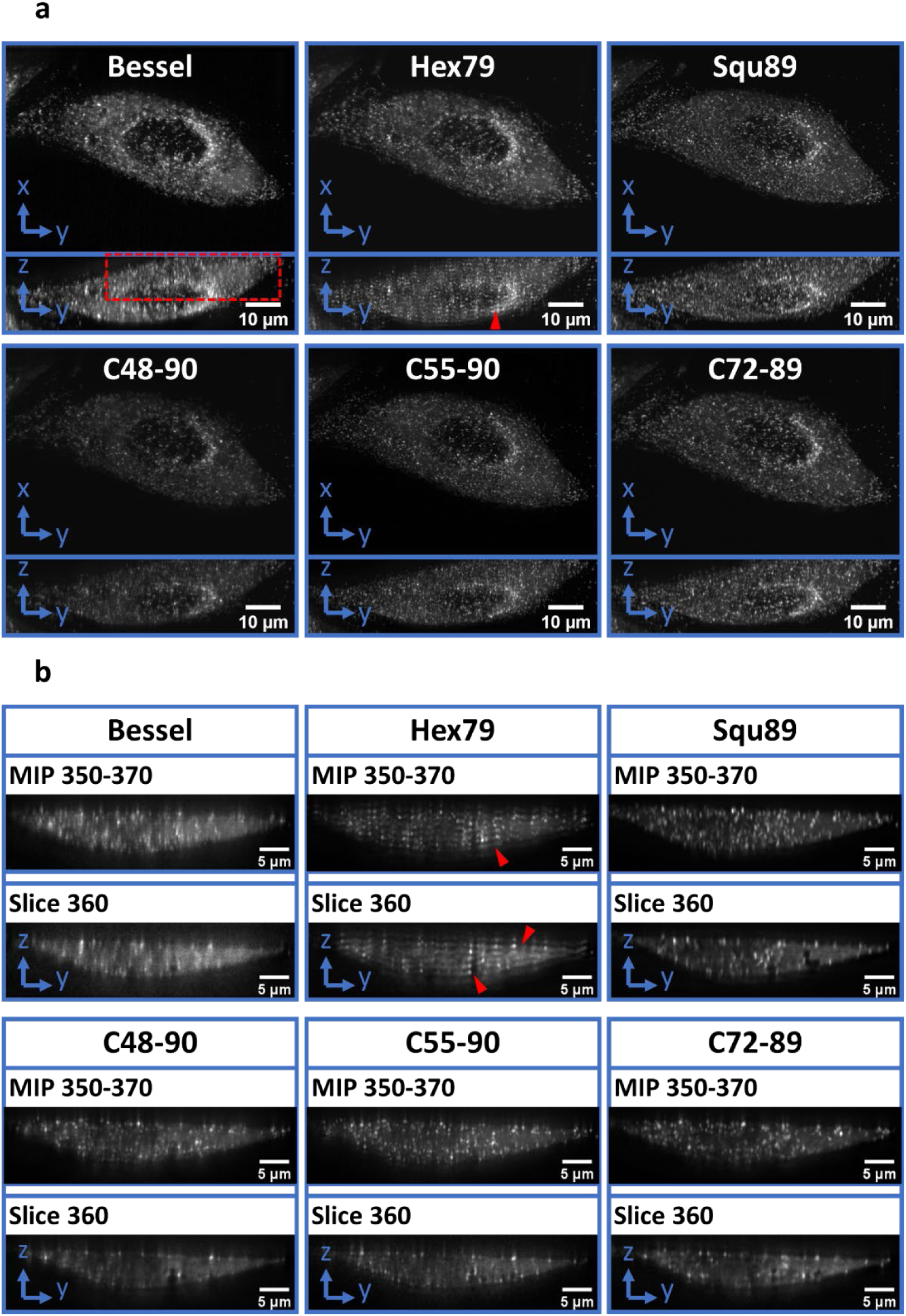
3D imaging of ARPE cells labeled with eGFP clathrin light-chain. **a.** Y-X and Y-Z maximum intensity projections of deconvolved image data as acquired by different light-sheets. **b.** Y-Z maximum intensity projections over a 2.1 microns range (top) and individual Y-Z sections from the red boxed region in a. Chevron marks deconvolution artifacts. MIP: maximum intensity projection. MIP range indicates depth in pixels (pixel size 104nm). Slices are centered at the middle of the MIP range.

## Discussion

We demonstrate flexible tuning of light-sheet thickness and confinement using Field synthesis. This has been achieved by scanning a beam only partially over an annular pupil filter, a procedure that we have termed C-light-sheets. The width of the main lobe of the light-sheet and the amount of out-of-focus excitation can be continuously tuned. This affords for much finer control over these two parameters compared to previous methods such as lattice light-sheet microscopy. In fact, of the two lattice light-sheet options (square and hexagonal), we found that only the more conservative square lattice has a generally usable optical transfer function. In contrast, the hexagonal lattice light-sheet exhibits gaps in its OTF, which make deconvolution much more demanding, and in our hands, results in artifacts in the reconstructed data.

Although we only demonstrated three C-light-sheets and two hexagonal light-sheets in this article, we have investigated more C-light-sheets and hexagonal light-sheets since it is very simple to generate different light-sheets with Field synthesis. We have generated hexagonal lattice light-sheets with even stronger sidelobes (>50% of the main lobe) but they were not useful for practical imaging. We have also generated more aggressive C-light-sheets by scanning a larger portion of the annular pupil filter, which produced a stronger beam skirt, which caused challenges in the deconvolution step. Obviously, a C-light-sheet with a large scanning range converges to a Bessel beam light-sheet.

We think that our new light-sheets will enable light-sheet users more options than the current state of the art to adjust their axial resolution and excitation confinement to their imaging applications. We also think that the Field Synthesis theorem has given a more intuitive approach to understand the characteristics of popular light-sheets such as Bessel and optical lattices. Thereby Field Synthesis has allowed us to engineer new light-sheets in a straightforward and rational way. In contrast, this would be difficult to realize in a lattice light-sheet microscope setup, as the number of useful optical lattices is limited. It is thus our hope that more useful light-sheets may be discovered with field synthesis, potentially by using more complex pupil filters.

## Acknowledgement

Reto Fiolka is grateful for support by the Cancer Prevention Institute of Texas (CPRIT grant RR160057) and the National Institute of Health (NCI grant R33 CA235254-01). The authors thank Dr. Tadamoto Isogai for the preparation of the collagen sample. The authors are grateful to Dr. Rosa E. Mino, who generated and validated the ARPE Cell line and to Dr. Madhura Bhave, who maintained the Cells in culture. The authors also thank Dr. Meghan Driscoll for the help of rotationally averaging the PSF, Dr. Jungsik Noh for the help of measuring light-sheets properties, Dr. Mark Kittisopikul for the comments on C-light-sheets, and Dr. Kevin M. Dean for a thorough reading of the manuscript and helpful comments.

## Notes

#### Summary of Updates

Formatting of manuscript was improved. Typos corrected.

## References

[1] J. Huisken, J. Swoger, F. Del Bene et al., “Optical Sectioning Deep Inside Live Embryos by Selective Plane Illumination Microscopy,” Science, 305(5686), 1007–1009 (2004).

[2] P. J. Keller, A. D. Schmidt, J. Wittbrodt et al., “Reconstruction of Zebrafish Early Embryonic Development by Scanned Light Sheet Microscopy,” Science, 322(5904), 1065–1069 (2008).

[3] B.-C. Chen, W. R. Legant, K. Wang et al., “Lattice light-sheet microscopy: Imaging molecules to embryos at high spatiotemporal resolution,” Science, 346(6208), (2014).

[4] Y. Wu, P. Wawrzusin, J. Senseney et al., “Spatially isotropic four-dimensional imaging with dual-view plane illumination microscopy,” Nature Biotechnology, 31, 1032 (2013).

[5] Kevin M. Dean, P. Roudot, Erik S. Welf et al., “Deconvolution-free Subcellular Imaging with Axially Swept Light Sheet Microscopy,” Biophysical Journal, 108(12), 2807–2815 (2015).

[6] Erik S. Welf, Meghan K. Driscoll, Kevin M. Dean et al., “Quantitative Multiscale Cell Imaging in Controlled 3D Microenvironments,” Developmental Cell, 36(4), 462–475 (2016).

[7] P. Theer, D. Dragneva, and M. Knop, “πSPIM: high NA high resolution isotropic light-sheet imaging in cell culture dishes,” Scientific Reports, 6, 32880 (2016).

[8] J. Durnin, J. J. Miceli, and J. H. Eberly, “Comparison of Bessel and Gaussian beams,” Optics Letters, 13(2), 79–80 (1988).

[9] G. A. Siviloglou, J. Broky, A. Dogariu et al., “Observation of Accelerating Airy Beams,” Physical Review Letters, 99(21), 213901 (2007).

[10] T. Vettenburg, H. I. C. Dalgarno, J. Nylk et al., “Light-sheet microscopy using an Airy beam,” Nature Methods, 11, 541 (2014).

[11] T. A. Planchon, L. Gao, D. E. Milkie et al., “Rapid three-dimensional isotropic imaging of living cells using Bessel beam plane illumination,” Nat Meth, 8(5), 417–423 (2011).

[12] F. O. Fahrbach, and A. Rohrbach, “Propagation stability of self-reconstructing Bessel beams enables contrast-enhanced imaging in thick media,” Nat Commun, 3, 632 (2012).

[13] F. O. Fahrbach, P. Simon, and A. Rohrbach, “Microscopy with self-reconstructing beams,” Nature Photonics, 4, 780 (2010).

[14] F. O. Fahrbach, and A. Rohrbach, “A line scanned light-sheet microscope with phase shaped self-reconstructing beams,” Optics Express, 18(23), 24229–24244 (2010).

[15] B.-J. Chang, M. Kittisopikul, K. M. Dean et al., “Universal light-sheet generation with field synthesis,” Nature Methods, (2019).

[16] T. Zhao, S. C. Lau, Y. Wang et al., “Multicolor 4D Fluorescence Microscopy using Ultrathin Bessel Light Sheets,” Scientific Reports, 6, 26159 (2016).

[17] B. Yang, X. Chen, Y. Wang et al., “Epi-illumination SPIM for volumetric imaging with high spatial-temporal resolution,” Nature Methods, 16(6), 501–504 (2019).

[18] B. Bailey, D. L. Farkas, D. L. Taylor et al., “Enhancement of axial resolution in fluorescence microscopy by standing-wave excitation,” Nature, 366(6450), 44–48 (1993).

[19] F. Lanni, B. Bailey, D. L. Farkas et al., “Excitation field synthesis as a means for obtaining enhanced axial resolution in fluorescence microscopes,” Bioimaging, 1(4), 187–196 (1993).

[20] M. G. Gustafsson, D. A. Agard, and J. W. Sedat, “I5M: 3D widefield light microscopy with better than 100 nm axial resolution,” J Microsc, 195(Pt 1), -16.

[21] S. Hell, and E. H. K. Stelzer, “Properties of a 4Pi confocal fluorescence microscope,” Journal of the Optical Society of America A, 9(12), 2159–2166 (1992).

[22] J. Bewersdorf, R. Schmidt, and S. W. Hell, “Comparison of I5M and 4Pi-microscopy,” Journal of Microscopy, 222(2), 105–117 (2006).

[23] M. Nagorni, and S. W. Hell, “Coherent use of opposing lenses for axial resolution increase in fluorescence microscopy. I. Comparative study of concepts,” Journal of the Optical Society of America A, 18(1), 36–48 (2001).

[24] S. W. Hell, S. Lindek, and E. H. K. Stelzer, “Enhancing the Axial Resolution in Far-field Light Microscopy: Two-photon 4Pi Confocal Fluorescence Microscopy,” Journal of Modern Optics, 41(4), 675–681 (1994).

[25] J. C. Gutiérrez-Vega, M. D. Iturbe-Castillo, and S. Chávez-Cerda, “Alternative formulation for invariant optical fields: Mathieu beams,” Optics Letters, 25(20), 1493–1495 (2000).

[26] F. O. Fahrbach, V. Gurchenkov, K. Alessandri et al., “Self-reconstructing sectioned Bessel beams offer submicron optical sectioning for large fields of view in light-sheet microscopy,” Optics Express, 21(9), 11425–11440 (2013).

[27] D. Loerke, M. Mettlen, D. Yarar et al., “Cargo and Dynamin Regulate Clathrin-Coated Pit Maturation,” PLOS Biology, 7(3), e1000057 (2009).

