## Supplementary Material for "Light-sheet engineering using the Field Synthesis theorem"

**Supplementary Note**

To synthesize a Lattice light-sheet (square or hexagonal) with Field Synthesis, we have previously stepped a line over three discrete scan positions in a pupil plane (corresponding to the main diffraction orders in the pupil in a conventional lattice light-sheet microscope)[15].

However, the central order (the line position at the center of the annulus) carries relatively little weight.

We thus simulated what difference it makes if one leaves the central order out. Surprisingly, there is relatively little change to the light-sheets profile, which makes us stipulate that the hexagonal lattice light-sheet is essentially a Cosine-Gauss beam.

For this manuscript, we thus left the central order away, both for the conventional lattices as well as the C-light-sheets.

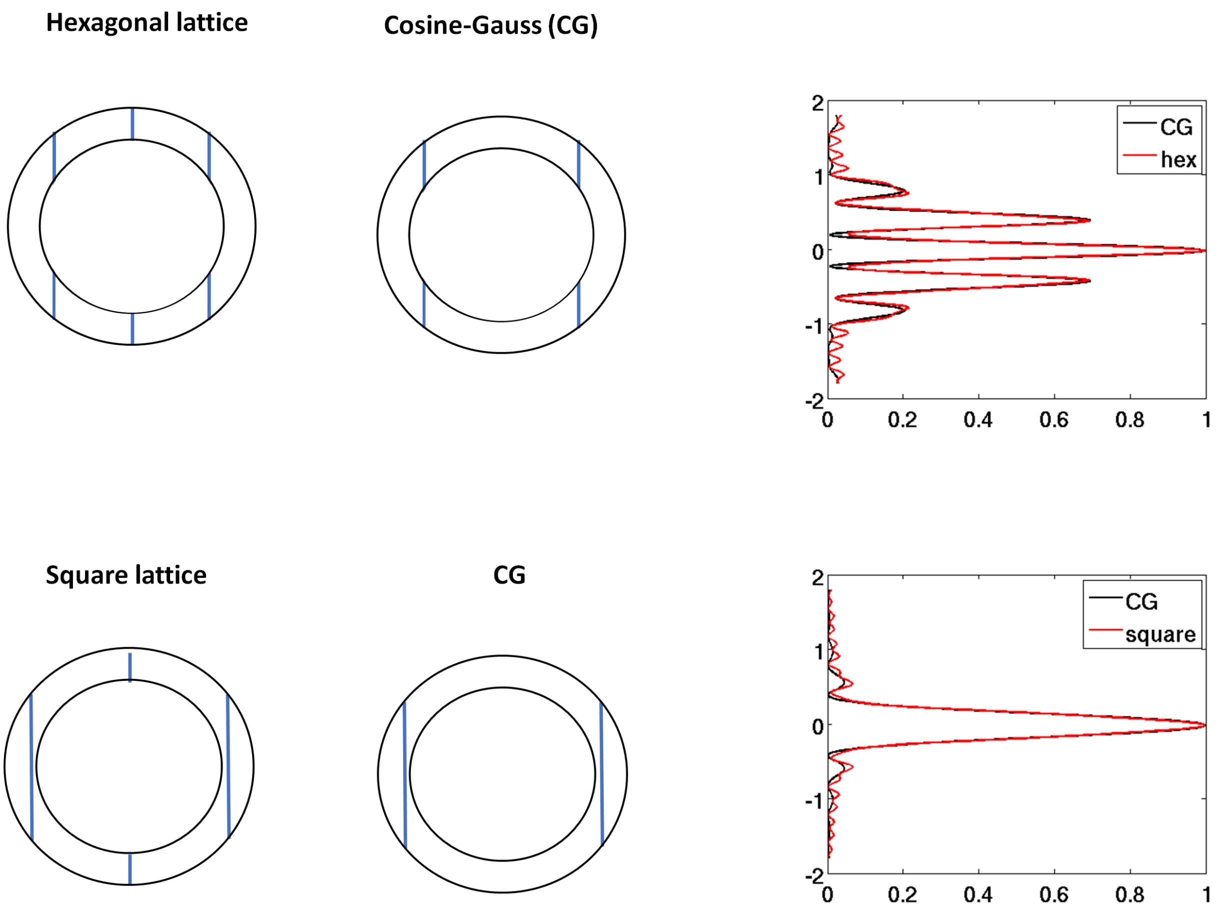


Numerical simulations of Lattice light-sheets generated by the Field Synthesis method, with (black curve in axial plot) and without (red curve in axial profile plot) the central order in the pupil. Pupil functions are shown on the left, the axial profiles through the center of the light-sheet are shown on the right.
